# Large-scale automated detection of gray whales off California in panchromatic and multispectral satellite imagery

**DOI:** 10.64898/2026.04.15.718679

**Authors:** Ludwig Houegnigan, Eduardo Cuesta Lazaró

## Abstract

Increasing human activities along the US west coast are of concern for populations of cetaceans and particularly for a number of large whale species that are recovering from overexploitation during the era of commercial whaling. New rapid monitoring tools, such as satellite imagery analysis powered by recent advances in artificial intelligence, have potential to provide additional broad-scale and near real-time capacities for survey and monitoring. This paper investigates and demonstrates the feasibility of automatic detection of gray whales in sub-meter satellite imagery off the coast of California, USA. Observations and statistical analysis of regional imagery allowed not only an assessment of their detectability but also the development of robust signal processing and machine learning-based solutions for automated detection. To that end, a regional dataset of 221 gray whales was created using signal processing to inform a deep-learning-based detection framework, and 20 different large neural network architectures for feature extraction followed by a support vector machine algorithm for classification were evaluated for their detection performance. Neural network backbones included 19 convolutional neural networks and 1 transformer network. The best architecture generally achieved satisfying performance with an average balanced accuracy reaching up to 99.90%. It is also demonstrated that panchromatic imagery, in spite of the lesser amount of information provided, can be used to perform detection with a relatively high accuracy of 87.05%, allowing wider spatial and temporal coverage. Large-scale deployment of the best performing models over a broad range of regional satellite imagery resulted in the detection of 3353 gray whales, as well as opportunistic detections of humpback, blue and fin whales, in and going from December 28th 2009 to March 26th 2023. It also provided meaningful data points concerning the migration routes of gray whales within the Channel Islands and Southern California Bight. The large number of high-confidence detections indicates the capacity for a large-scale monitoring approach to support state and federal conservation policies such as gear mitigation, vessel speed reduction programs, or shipping lane redefinition that could also be expanded to other areas and for other species.

## 1. Introduction

While many whale populations are recovering from historical impacts of commercial whaling, ongoing human activities in the marine environment, such as shipping, fishing, noise and other sources of disturbance, remain threats to whale populations worldwide [30, 31, 32]. Climate change furthermore creates new threats to whales through ecological disruptions that can cause whales to shift foraging and migration patterns and timing [38–40].

These impacts to large whales represent a growing concern off the coast of California, which features key migratory routes and feeding grounds for populations of gray whales (*Eschrichtius robustus*), humpback whales (*Megaptera novaeangliae*), blue whales (*Balaenoptera musculus*) and fin whales (*Balaenoptera physalus*). These migratory routes and feeding grounds seasonally overlap with busy shipping lanes, with particularly high risk of collision in the Santa Barbara Channel and approaching the Port of Long Beach as well as along the shipping lanes off San Francisco[29]. Furthermore, whale entanglement in fishing gear has increased in recent years, particularly impacting gray and humpback whales (with 208 and 165 entanglements reported between 1982 and 2017, respectively) but also affecting blue and fin whales [2]. Ship strikes are estimated to impact whales at high rates as well, including suggestions that they are the major factor in limiting blue whale recovery [41,42]. All marine mammals in US waters are protected under the Marine Mammal Protection Act (MMPA), while humpback, blue, fin and western North Pacific gray whales found off the California coast are also listed as endangered under the US Endangered Species Act (ESA) [1,34].

Significant monitoring, management, research, and mitigation efforts have been established to address these threats to whales. Key interventions include traffic separation schemes to reduce the co-occurrence of whales with commercial shipping, voluntary vessel speed reduction programs, and the designation of seasonal whale advisory zones. These programs rely on data from a variety of monitoring initiatives such as Whale Safe that integrates systematic and opportunistic sightings, passive acoustic detection and model output to understand and monitor large whale migration and distribution patterns and to inform whale population assessments [4,5,6,7]. Monitoring data from traditional vessel-based or aerial surveys can also be leveraged in combination with oceanographic data to inform development and validation of species habitat models [42]. However, consistent deployment of traditional methods such as vessel-based or aerial surveys can be cost prohibitive and logistically challenging [8,9]. As a result, substantial gaps persist in our understanding of the variability in large whale distribution patterns. Near real-time data is therefore needed to fulfill management requirements to protect whales from anthropogenic threats globally and in California specifically.

The use of sub-meter satellite imagery as a complementary monitoring tool for wildlife has gained traction in recent years due to its unique capability to image large areas without the logistical complexity of conventional *in situ* survey tools. Wildlife megafauna such as large whales can be observed and detected in sub-meter resolution satellite imagery. The number of studies estimating the number of individuals of different species (polar bears, seals, right whales, etc) or estimating population size from sub-meter imagery is constantly increasing. However, a large number of studies rely on manual review of imagery for animal detections; this is a valuable and unavoidable step in the development of annotated datasets for automation, yet it can also be a slow and tedious process because a single multispectral image tile covering only 25 km^2^ can, depending on resolution, require the analysis of over 2 billion pixels. For satellite imagery to be operationalized for use in science and conservation decision-making, faster and more reliable detection tools must be developed to streamline the review process over large spatiotemporal scales.

Progress in machine and deep learning over the past two decades has contributed to significant performance improvement in a wide range of fields such as natural language processing, speech processing and image processing [10]. A central tenet to this progress was the existence of well-defined problems with large annotated databases that were typically built through decades of research and frequently contained millions of statistically relevant samples, without which the training of such algorithms generally results in suboptimal performance. The use of deep learning approaches for the detection of marine mammals in satellite imagery is a relatively new development that at this point is supported by non-standardized and small datasets; achieving high accuracy results in this context is not a straightforward endeavor.

For the automated detection of whales in satellite imagery, a key framework was proposed by Guirado [43] which consisted first in (1) dividing the image into cells and in classifying each cell according to its probability of containing a whale and then (2) in applying an object detector (in that case Faster-RCNN) to pinpoint the position of whales within cells and to produce an exact count. In such an approach, classification is the key step while detection (in some cases referred to as localization) produces a precise location within the cell and a refined count in case multiple animals are present per cell. Multiple approaches are available to address both (1) and (2). Other research efforts focused directly on object detection using a state of the art algorithm such as YOLO [37]. In those cases classification and object detection performance have typically been hindered firstly by a small database which produces suboptimally trained networks and secondly by using only the RGB bands, the combination of those two factors have potential to produce a large number of false positives and negatives which are inadequate for automated systems for broad-scale geospatial datasets.

This paper undertakes classification of cells using a different approach that is more tailored to smaller datasets (in the context of CNN-based machine learning, datasets are considered here to be “small” when they contain below 10000 samples, even datasets with multiple of thousands of whales should be considered “small” in the context of machine learning) and aims to provide high accuracy at classification so as to minimize false positives and negatives. Classification is optimized by testing a large number of feature extraction schemes and by using all available frequency bands so as to extract a maximum of information from data. This is done by extracting features with pretrained convolutional and then using a support vector machine algorithm to classify said features.

### 1.1. Focal species and study area

Gray whales (*Eschrichtius robustus*) undertake one of the longest migrations of any mammal species at over 20,000 km[11,12]. Gray whales are therefore considered as “sentinels of climate change” [26,27]: their summer distribution in their Arctic feeding grounds is strongly related to the local and seasonal increase in food abundance and to the potential opening of new opportunistic migration routes due to increased warming and melting ice areas. In the Eastern Pacific, gray whales typically migrate from their wintering and nursery ground in Baja California, Mexico, up to their summering ground in the northernmost locations of Alaska past the Bering Strait. Detailed counts are performed each year at key locations along this well-documented migration route [13,14]. The population of eastern North Pacific gray whales has recovered from the brink of extinction due to commercial whaling in the early 20th century to an estimated abundance of 16,650 in 2022[16], with a current conservation status assessed as ‘’least concern’[28]. However, the current population has experienced a rapid and concerning decrease from an estimated 26,960 in 2016 to 20,580 in 2020[16] and an Unusual Mortality Event (UME) was declared in 2019[16]. For the population of western North Pacific gray whales, listed as endangered under the Endangered Species Act with an estimated number under 300 individuals, it was also found that some individuals perform an even larger migration to winter in the lagoons of Baja California where they share habitat with the eastern gray whale population[15,17], which implies that monitoring and conservation efforts put in place along the Eastern Pacific migration route could have a positive impact for both populations.

Once common throughout the Northern Hemisphere, the gray whale was extinct in the Atlantic by the early 1700s [57] but recent sightings have been reported of lone gray whales in the waters of Spain (2010, 2021, 2023), Israel (2010), Namibia (2013), and France (2023) were reported [18] and are hypothesized to be related to the ephemeral opening of the Canadian Northwestern passage resulting from sea ice retreat driven by climate change.. An occurrence in Hawaii (2022) was also reported [19]. Therefore, the ability to track gray whales in areas where they have long been absent may also shed light on the influence of climate and ecosystem changes on marine wildlife occurrence and distribution.

While advances in machine learning approaches can reduce the quantity of data required for high performance supervised learning methods [44,45], a relatively large amount of data is still needed to train efficient deep learning and transfer learning methods. Therefore, well-studied and relatively abundant whale populations can serve as a valuable case study for automated detection. This is the case for eastern North Pacific gray whales (*Eschrichtius robustus*) whose migration route from wintering areas in Baja California, Mexico, up to the Bering and Chukchi seas is well-studied. In Laguna Ojo de Liebre, Laguna San Ignacio or Bahia Magdalena, aggregations of multiple hundreds to multiple thousands of gray whales can be observed on certain days within a few tens of kilometers. Furthermore, during their migration, eastern gray whales stay relatively close to shore, where significantly more satellite data is available than in the high seas making them adequate candidates for detection in satellite imagery and for the training of a successful recognition system. Shore-based counts are performed each year at multiple locations off the coast of California, such as Yaquina Head, Granite Canyon, Diablo Canyon and Point Vicente, thus documenting the timing and abundance of their migration [13,14]. Wintering grounds often feature calm, clear waters and large persistent aggregations of whales offering readily available access to study gray whales via satellite imagery, but may not generalize to other environments. Coastal California presents a unique test case for developing a whale detection algorithm able to perform across a broad spectrum of water color, surface roughness turbidity levels, a diversity and large amount of submerged or floating objects that could trigger false detections that could not always be observed in Baja California (kelp, foam, floating debris, etc), and a diversity of marine wildlife including eight species of large whales. Thus, detecting gray whales in open waters along the California coast, beyond the more sheltered wintering lagoons, offers a unique test-case with a realistic indication of the performance that can be expected of a whale detection system.

## METHODS

### 2. Data processing and analysis

#### 2.1 Satellite imagery preprocessing

Panchromatic and multispectral imagery were both utilized for this study. Multispectral data comprised imagery from Maxar’s Worldview 2 (WV2) and Worldview 3 (WV3), while panchromatic imagery derived from Maxar’s Worldview 1 (WV1). WV2 and WV3 provide 8 multispectral bands; the use of WV1 which features a single panchromatic band provides additional spatiotemporal coverage and therefore could support better description of migratory distributions and density estimates, provided that acceptable detection rates can be obtained. Examples of Gray whales in WV1, WV2 and WV3 imagery are presented with the corresponding estimated number of pixels and ground sampling distance are presented in Tables 2.1 to 2.3. Lower amounts of pixels induced by a lower ground sampling distance will imply a smaller cross sectional area of the target which will typically result in lower detection performance [20, 21].

Prior to detection analysis, orthorectification was performed for all coastal images and top-of-atmosphere reflectance values were calculated to account for sensor exposure and sun irradiance changes and taking into account geospatial parameters (sun elevation angle, earth-sun distance at time of acquisition, etc). Pansharpening was then applied using a standard IHS algorithm [22] to bring the lower resolution multispectral bands (1.24m at nadir) to the spatial resolution of the panchromatic band (0.31m at nadir).

##### 2.1.1 Extraction and analysis of whale pixels

**Figure 2.1.**
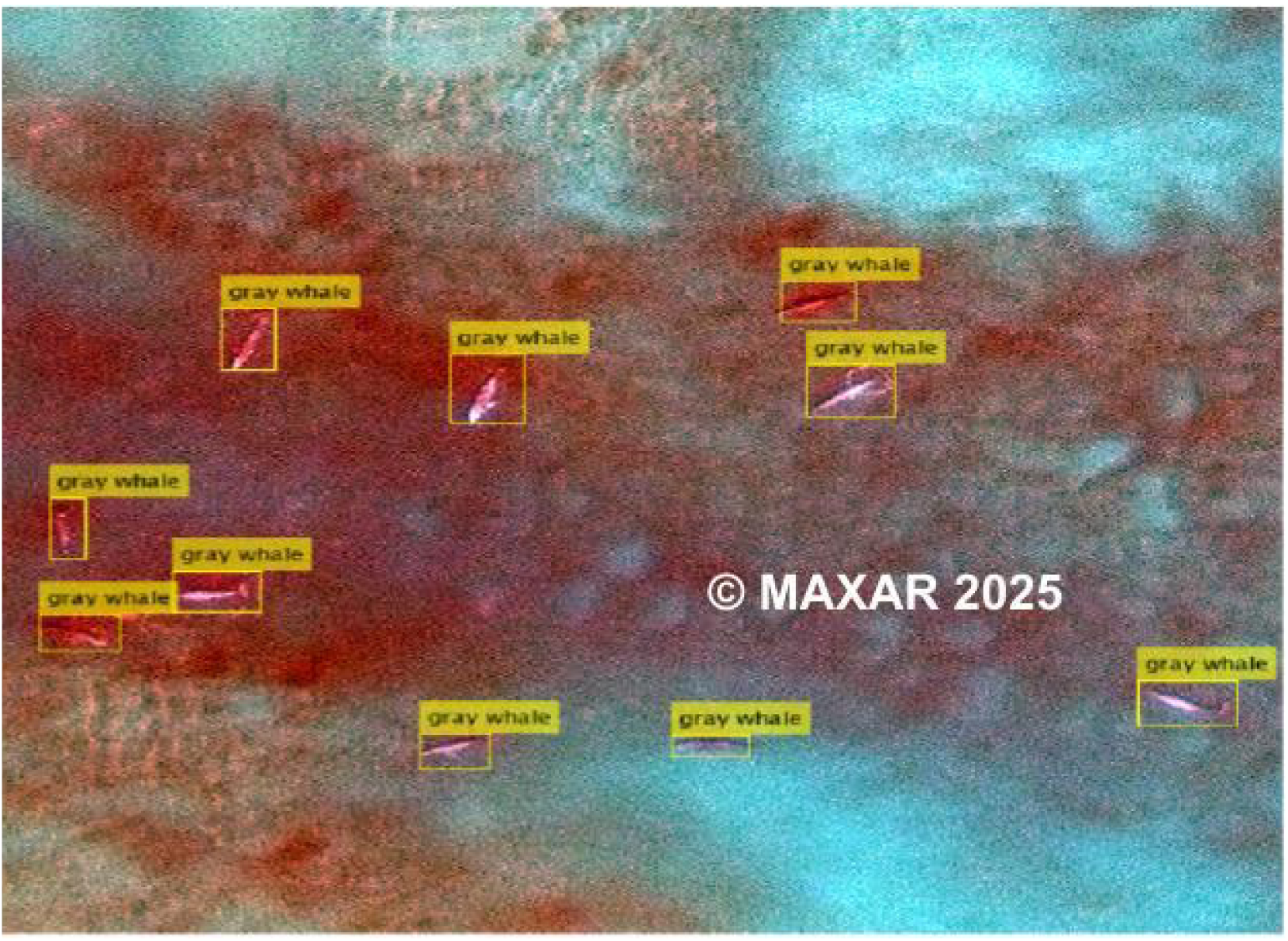
Example of aggregation of Gray whales observed in WV3 imagery at Boca de Santo Domingo, north of the Bahia Magdalena lagoon complex, Baja California, Mexico. This statistically contrasted image gives gray whales an artificial pink color tone.

Four satellite images over two wintering lagoons of the Baja California peninsula (table 2.4) were analyzed for pixel extraction and used to develop a predetection step outlined in section 2.2. A prerequisite to predetection was the extraction of pixels corresponding to the target of interest. To automatically extract pixel value and shape information from gray whale observations of interest, a series of classical morphological operations were implemented, as described in a previous publication [23]. This allowed us to visually examine and to quantitatively assess the pixel distribution for both gray whales and the surrounding marine environment for each of the multispectral bands through histograms. These showed that the higher frequency range of the color distribution (coastal, blue, green bands) offered better separation capabilities from the surrounding ocean colors, and then, as frequency decreased, separation between the whale pixel distribution and the water pixel distribution also decreased (see Figures SI- 2.1(a) to (h)). Therefore, the lower range of the frequency spectrum seemed to include less information for direct separation between water surface and gray whales.

**Table 2.4.**
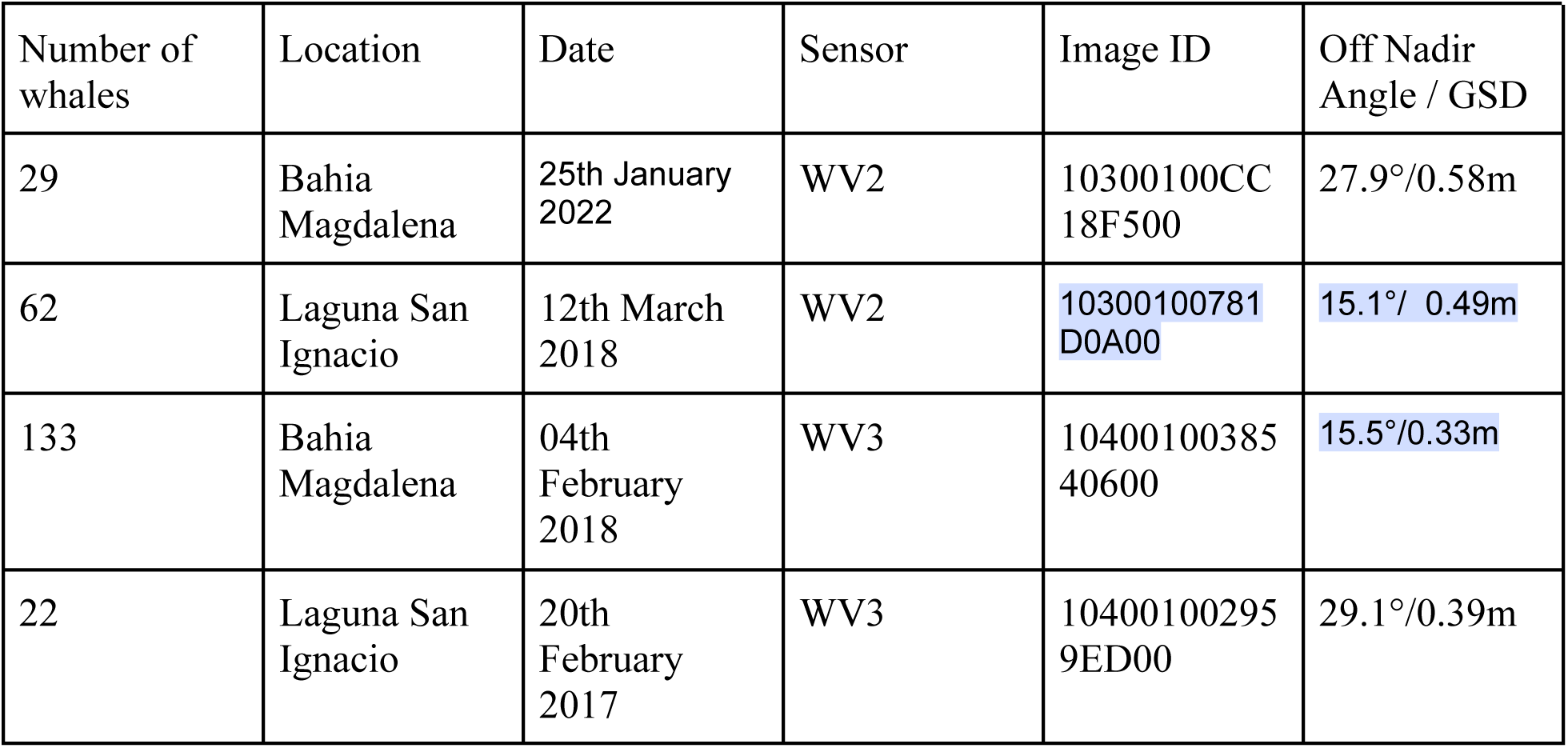
Satellite imagery location used in preprocessing.

This separation was further estimated numerically using firstly the Kullback-Leibler divergence (KLD) and secondly the estimated overlap between the gray whale pixel distribution and the water pixel distribution derived from histograms. The KLD is a measure of the dissimilarity between two distributions, higher values of the KLD indicating better separated distributions. KLD values were found to be higher for the coastal, blue and green bands which confirmed numerically the visual assessment of histograms (Table 2.1). The estimated overlap between distributions is also lower for the coastal, blue and green bands even though the estimate for the NIR2 band was found to be also relatively low. Whales were typically submerged near the ocean surface, therefore the extracted whale pixels are a mixture of the true whale pixel color and water pixel color. Even still, KLD estimates provided us with useful information to identify candidate pixels for the development of a whale detection algorithm.

**Table 2.1.**
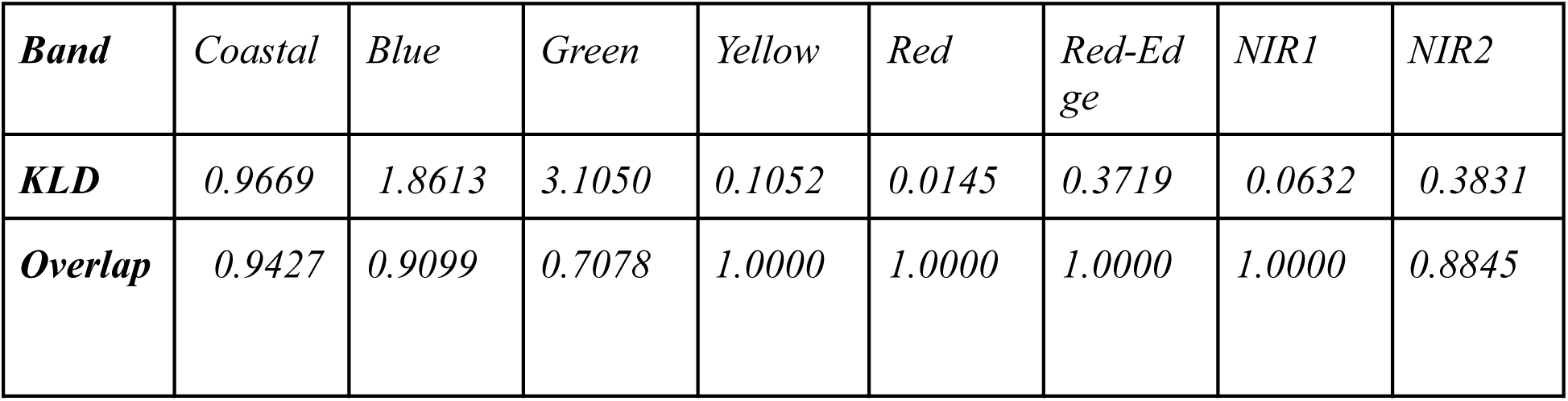
Kullback-Leibler divergence and overlap estimates per band describing the separation between gray whale and water pixel distribution (for WV2 and WV3 data).

| ***Band*** | *Coastal* | *Blue* | *Green* | *Yellow* | *Red* | *Red-Edge* | *NIR1* | *NIR2* |
| --- | --- | --- | --- | --- | --- | --- | --- | --- |
| ***KLD*** | *0.9669* | *1.8613* | *3.1050* | *0.1052* | *0.0145* | *0.3719* | *0.0632* | *0.3831* |
| ***Overlap*** | *0.9427* | *0.9099* | *0.7078* | *1.0000* | *1.0000* | *1.0000* | *1.0000* | *0.8845* |

### 2.2 Predetection and dataset building using signal processing

Subspace methods typically used in signal processing for frequency estimation as well as for source localization problems [24] offer a simple and robust solution to perform rapid detection with a small sample support where deep-learning based methods typically require more samples [25]. However those methods, in their basic form, focus on multispectral intensity values and do not take into account spatial and shape information. This predetection step using subspace methods allowed to rapidly review imagery and to generate a regional dataset of California gray whales. All extracted whale multispectral samples obtained in the breeding grounds were used to define two complementary subspaces: (1) a gray whale signal subspace, and (2) a noise subspace. The gray whale signal subspace was defined using 220 gray whales that were identified in Baja California imagery data (eq.1). The noise subspace (eq.2) was then defined by selecting pixels in regions of the image which contained only the ocean surface and were void of whales or any other object. Figure 2.2(a) illustrates a natural color image with 2 clearly identified gray whales and Figure 2.2(b) illustrates the detection response output of a subspace-based method.

The whale signal subspace and noise subspace used in this development were defined as follows:

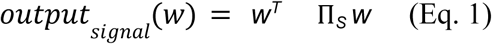

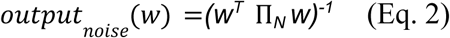

*Where:*

w is the input pixel vector

Π*_S_ is the signal subspace matrix, built with the signal eigenvectors of the covariance matrix of whale pixels[55,56]*

Π*_N_ is the noise subspace matrix, built with the noise eigenvectors of the covariance matrix of water surface pixels*.

**Fig. 2.2.**
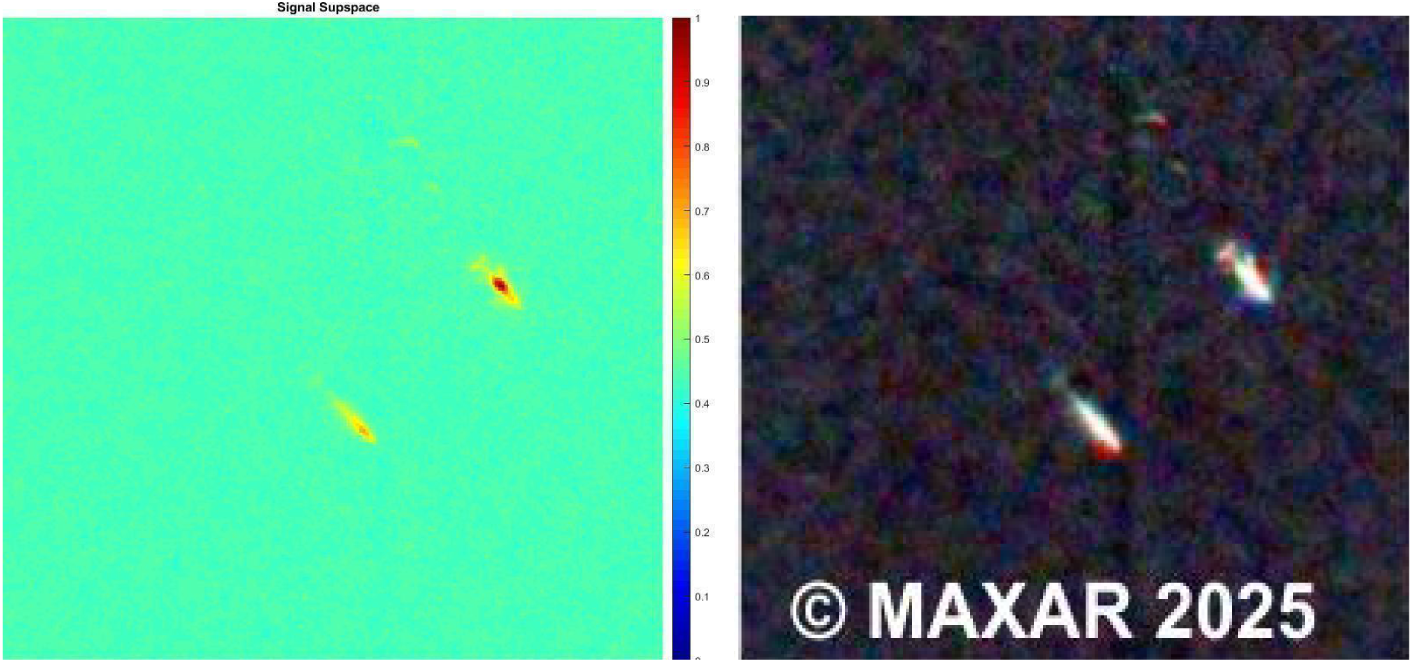
a (left) and Fig 2.2b (right). Figure 2.2 illustrates the subspace-based response output for two gray whales that are clearly identifiable in the corresponding natural color image.

The output of this process was reviewed and allowed the detection of an initial dataset of 221 gray whale detections within the California region. Those imagery samples were subsequently included in a California regional training dataset which was then used to train a high-accuracy automated detection system for the coastal California region. Samples of gray whales in addition to other items detected through the subspace algorithm are presented in the Supplemental Information section 3.2.

### 3. Detection methodology

#### 3.1 Feature extraction using pretrained convolutional neural networks and support vector machines

Subspace methods allowed the rapid pre-detection of gray whales based on multispectral intensity values using very few samples of targets of interest. Extracting features from pretrained CNNs offered a more robust detection approach harnessing the high performance and discriminative capability of deep neural networks pretrained on large datasets with more than a million RGB images such as Imagenet or Places 365, and to transfer those abilities to a less well-defined problem with a smaller sample support, such as the current gray whale detection problem [33]. However, the main limitation of pretrained networks is that their dimensions are scaled for RGB inputs from natural images and are not meant for multispectral imagery that typically has over 4 bands. To tackle this limitation, multiple options were investigated in this study: (1) dimensionality reduction using principal component analysis, (2) direct use of the RGB channels and (3) use of the full multispectral content.

Using one of the final layers of a CNN, robust and general image features can be extracted. Then a Support Vector Machine (SVM) can be trained on labeled extracted features with results that are typically better than training from scratch or than fine-tuning a CNN, particularly when the sample support for training is small (see Supplemental Information section 3.1) for further processing details). For the purpose of our recognition task, 20 different CNNs were used for feature extraction and the extracted features from each were then fed to a support vector machine architecture for classification.

#### 3.2 Detection problem formulation

For simplicity, detection was formulated as a classification problem involving five different classes. The gray whales class (1) was composed of 221 multispectral gray whale samples from California imagery, from which 121 were used for training and validation and 100 were used for testing. The “ships and container ships” class (2) was composed of 95 multispectral ships and container ships from California imagery, from which 52 were used for training and validation and 43 were used for testing Figures SI-3.5(a-c). The “planes and helicopters” class (3) was composed of 89 multispectral planes and helicopters from California imagery, from which 52 were used for training and validation and 43 for testing. The “non-whale detectables” class (4) comprised 80 multispectral non-whale detectables from California imagery, from which 44 were used for training and validation and 36 for testing. Non-whale detectables (abbreviated as “nwd”) represent events of interest, seemingly on the surface or below the surface of water, that are not whales but have potential to confuse the whale detection/classification process. The “background class” (5) comprised 225 multispectral water surface samples with different water conditions from California imagery, from which 123 were used for training and validation and 102 for testing. Examples of backgrounds are shown in Fig 3.15(a) to Fig 3.15(f).

Cross validation was applied using all class samples to produce 100 randomly generated training and testing datasets. Each of the randomly generated datasets is then used for feature extraction and then fed to a support vector machine algorithm. Therefore, for each deep learning model used for feature extraction 100 SVMs are trained and the results are then averaged across cross-validation datasets.

Preprocessing divided a larger image into smaller cells of dimension 224 x 224 and assigned a class to each cell. Each cell was then used as input to the detection model and a cell was classified as positive if at least one whale was present within the cell and negative if no whale was present independently of the identification of the class (non-whale detectable, ship, background). Accuracy (ACC), True positive rate (TPR), True negative rate (TNR), False positive rate (FPR), and False negative rate (FNR) were used as metrics to assess the performance of the models. The number of False positives and False negatives per square kilometers were also estimated.

## 4. Results

Detection with multispectral imagery outperformed both RGB and panchromatic detections (Table 4.1.1) with respective accuracies of 99.9%%, 99.39%, and 87.05%. While those results seem relatively close to each other and all seem relatively high, they result in quite different false positive and false negative rates (Table 4.1.1). Compared to the 8-band multispectral, the panchromatic or RGB inputs represented a significant loss of information which reduced detection capability by decreasing the signal-to-noise ratio and increasing variance: for the panchromatic case, accuracy varied within a range of 42.55% (with a standard deviation σ=7.75%), 3.17% (σ= 0.76%) for RGB, and 2.2% (σ=0.56%) for 8-band multispectral. These results indicate that as the amount of information increases, the performance of networks is increasingly similar, whereas when minimal spectral information is available it becomes necessary to test a large amount of network architecture to optimize results. The observed improvements in variance resulting from additional information available in the 8-band multispectral is key towards achieving a fully automated reliable system as opposed to a system that would require human supervision (see SI for further information on detection outcomes and misclassifications at different spectral band combinations).

**Table 4.1.1.**
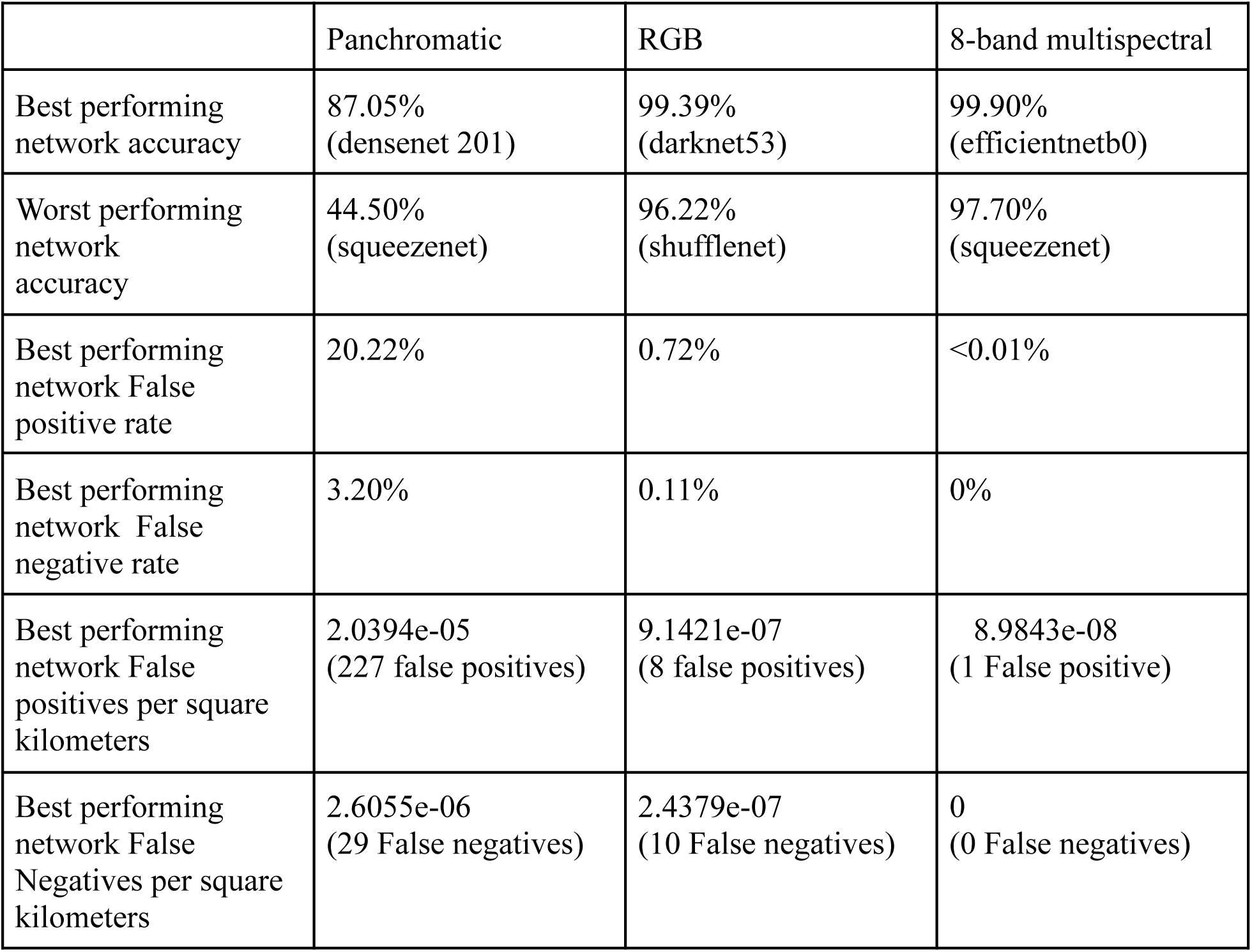
General results for detection for Panchromatic, RGB and 8-band multispectral input. In the first two rows, neural network architectures corresponding to each performance metric are indicated between brackets.

### 4.1 Misclassifications of 8-band multispectral inputs

The best multispectral classifier only produced one false positive (out of 324 test samples at each iteration), and that false positive appears to correspond to foam at the ocean surface. The use of multispectral data seems to resolve many of the complex situations RGB classifiers could not classify correctly, highlighting the discriminating capacity of multispectral inputs. The addition of higher frequency bands (i.e. coastal) and lower frequency (e.g. yellow to nir2) improves classification by a few tenth of a percent which significantly reduces the amount of false positives and false negatives (Fig 4.3(b)).

**Fig 4.3.**
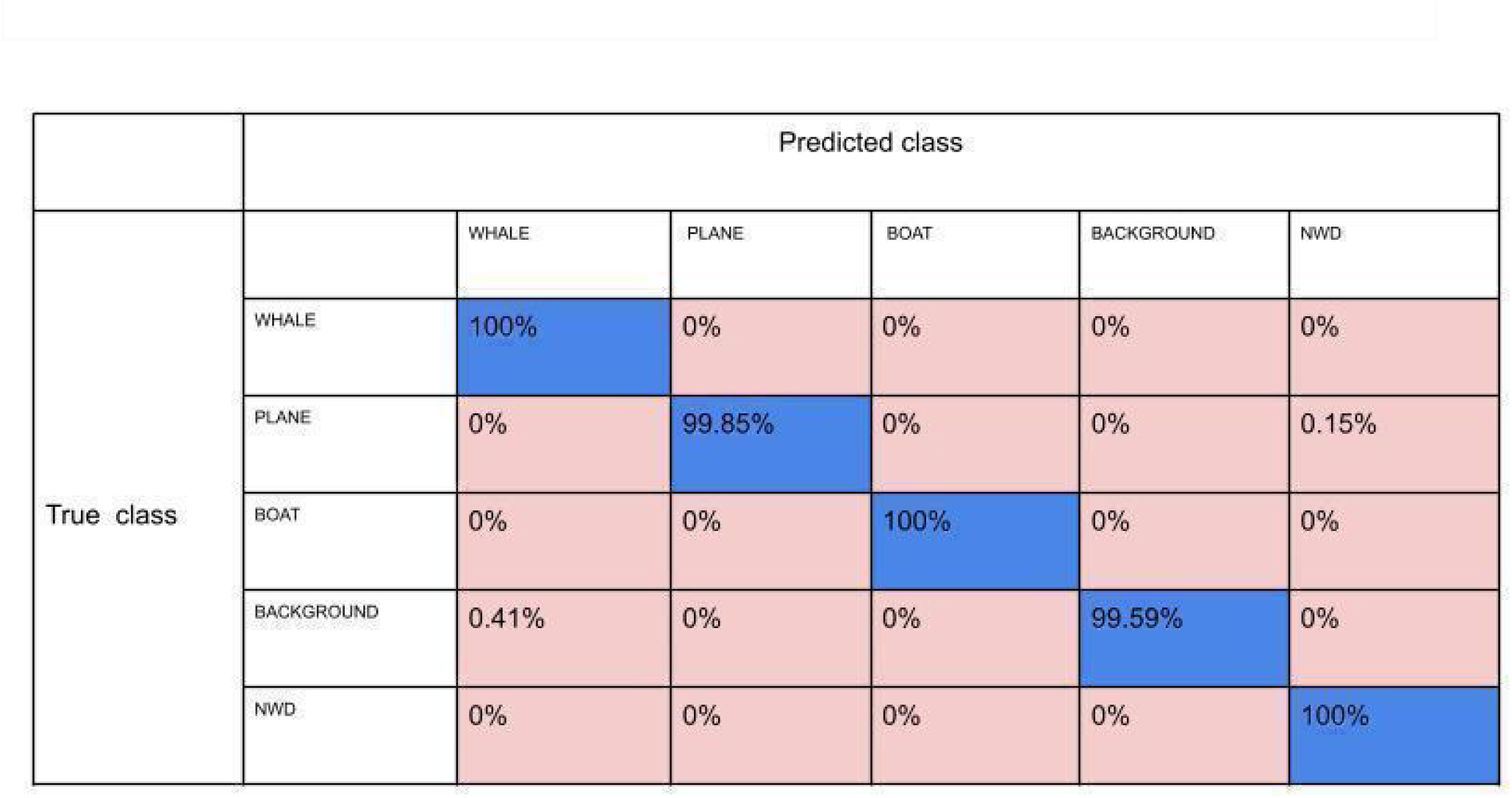
(b) confusion matrix for multispectral input

### 4.2 Large-scale deployment

The best performing models were deployed on approximately 624,000 square kilometers of WV2 and WV3 coastal California imagery. Initial emphasis was put on Southern California which offers lower turbidity and a lesser cloud coverage, particularly to an area spanning San Diego to Coronado Island, as well as another area near Granite Canyon where Gray whale surveys are performed yearly by NOAA [35,36]. Later imagery additions added coverage of nearly the entire coast of California as described in figures 4.2.1 and 4.2.2, showing the density of survey effort.

**Figure 4.2.1.**
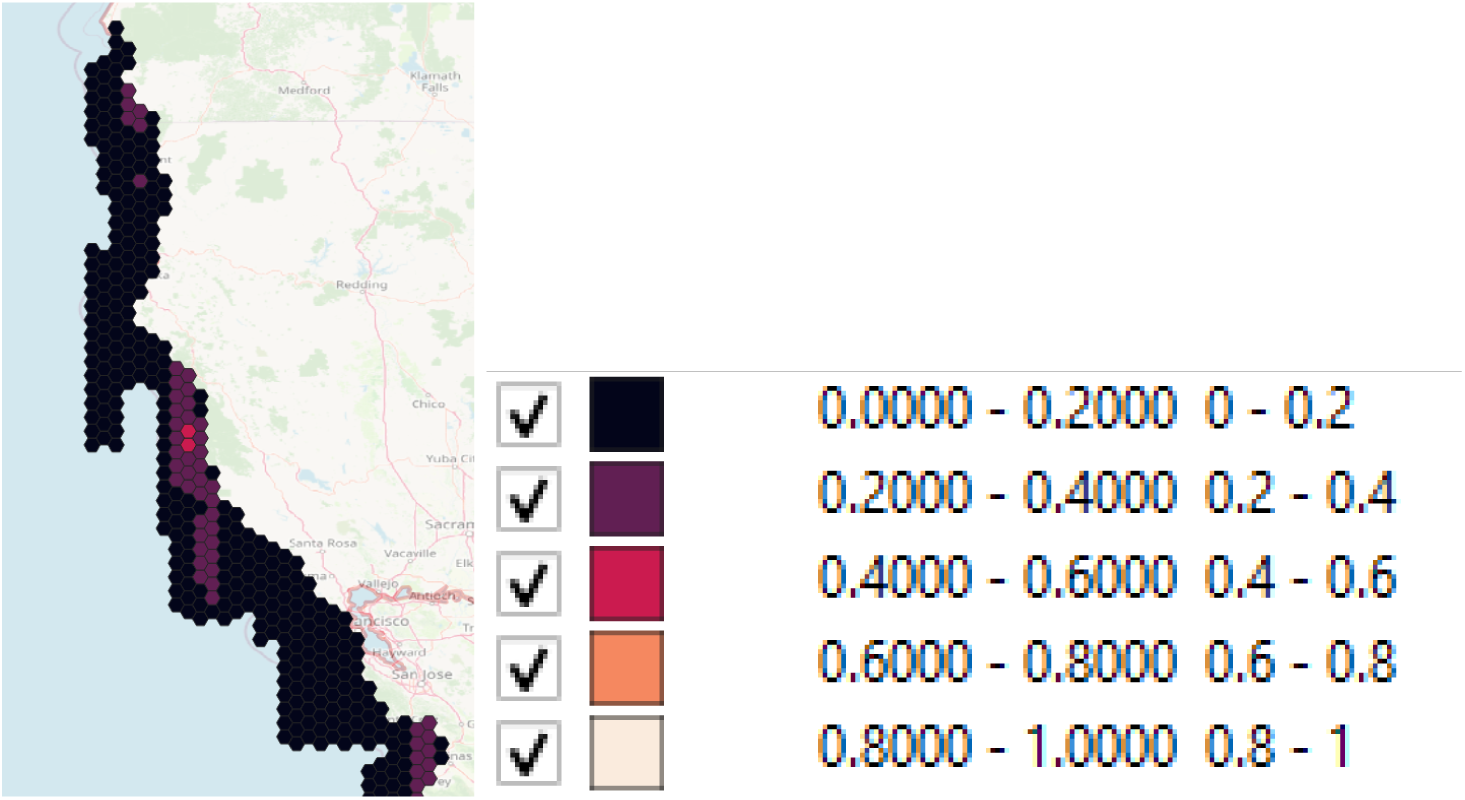
Northern portion of California survey effort (click image and follow link to zoom in)

**Figure 4.2.2.**
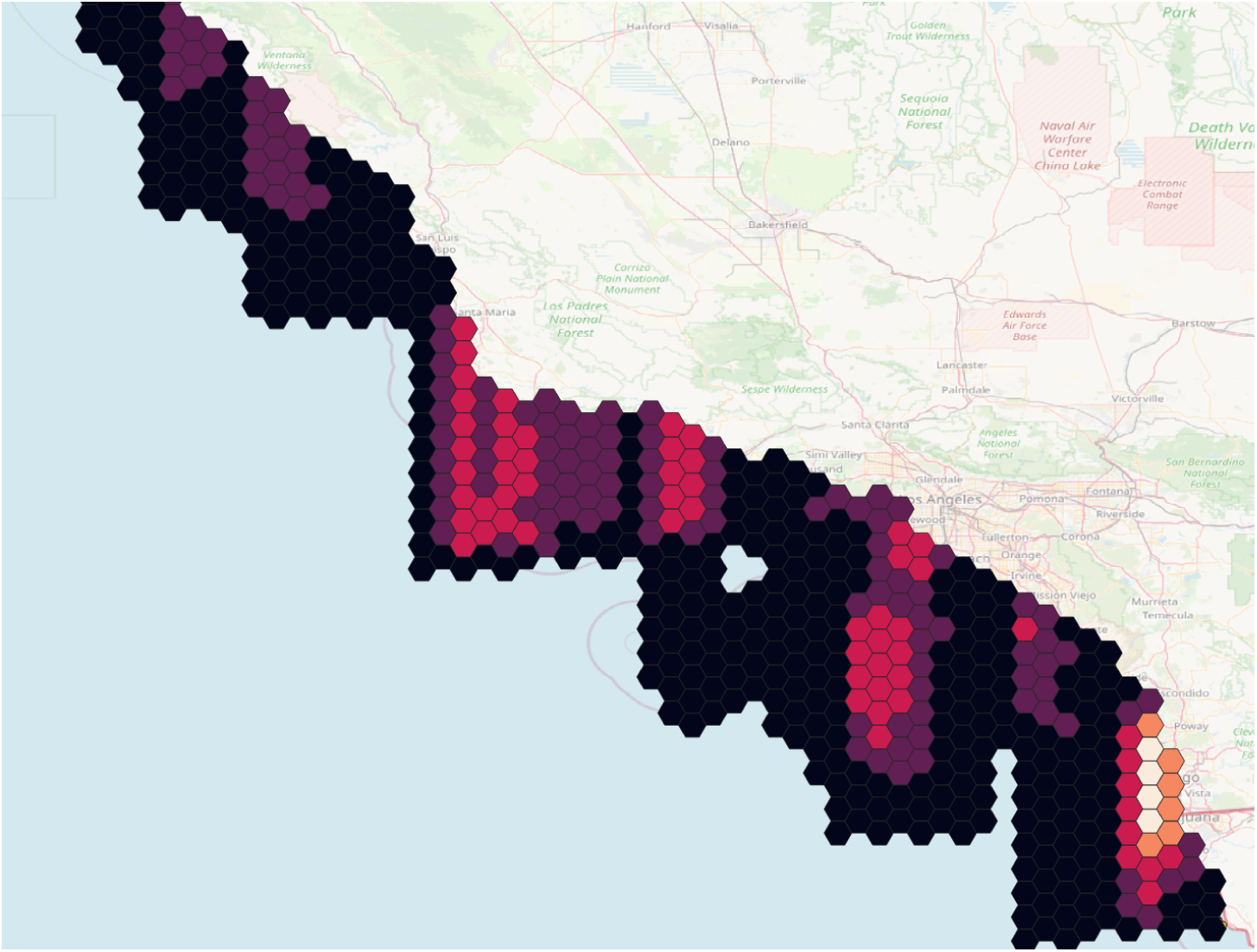
Southern portion of California survey effort (click image and follow link to zoom in)

3353 gray whales were detected over a time span going from December 28th 2009 (6 detections near Mattole Canyon State Marine Reserve) to March 26th 2023 (9 detections within Monterey Bay) and covering nearly the entire coast of California with northernmost detections near Crescent City (-112.0936°, 41.5345°) and a large number of detections near the US-Mexico border. The longitudinal coastal configuration near San Diego and the US-Mexico border favors the capture of archival imagery covering a large portion of ocean as opposed to the latitudinal coastal configuration north of the Channel Islands from Point Conception to Los Angeles. The earliest detections made for a given year was made on November 13th, 2015, near San Nicolas Island.

The overall median distance to the continental coast was found to be approximately 5.86 km (see Table 4.2.1) however this varied substantially between the Southern California Bight and the Northern California coastline (above San Francisco). Detections made closest to a coastline were made east and west of San Clemente Island, many of which were within 500 meters from the coastline, with the closest detection falling 50 meters from the coast. Detections furthest away from the coast were made between Santa Cruz Island and San Nicolas Island with a peak value at 57.2 km, 52 other detections were over 35 kilometers.

Observations made within the Santa Barbara Channel and the California Channel Islands confirm the existence of multiple migratory paths. This includes a well-known inshore route, as well as multiple less-studied offshore routes between the Northern Channel Islands (particularly between San Miguel, Santa Rosa, and Santa Cruz Islands) as well as near Santa Catalina and San Clemente Islands [12]. This study illuminates significant usage of habitats as far west as San Nicolas Island highlighting the importance of offshore migratory corridors traversing the southern California Bight and in particular along the west coast of San Clemente Island, an area frequently exposed to naval sonar within the Southern California Anti-Submarine Warfare Range (SOAR). A potential preferential use of offshore corridors could indicate avoidance of interaction with human activities (e.g. commercial shipping lanes) prevalent in the nearshore migration corridor but it could also correspond to a least cost path to traverse the southern California Bight. A more comprehensive description of migration through the Channel Islands will require the analysis of additional imagery in future work and the development of reliable density estimates (i.e. minimizing bias potentially introduced by sea state or cloud cover).

**Table 4.2.1.** Statistical measurements of distance to continental coastline.

| Percentile | 2.5 | 25 | 50 | 75 | 97.5 |
| --- | --- | --- | --- | --- | --- |
| Distance to coast | 0.86 | 2.85 | 5.86 | 11.50 | 30.37 |

**Figure 4.2.3.**
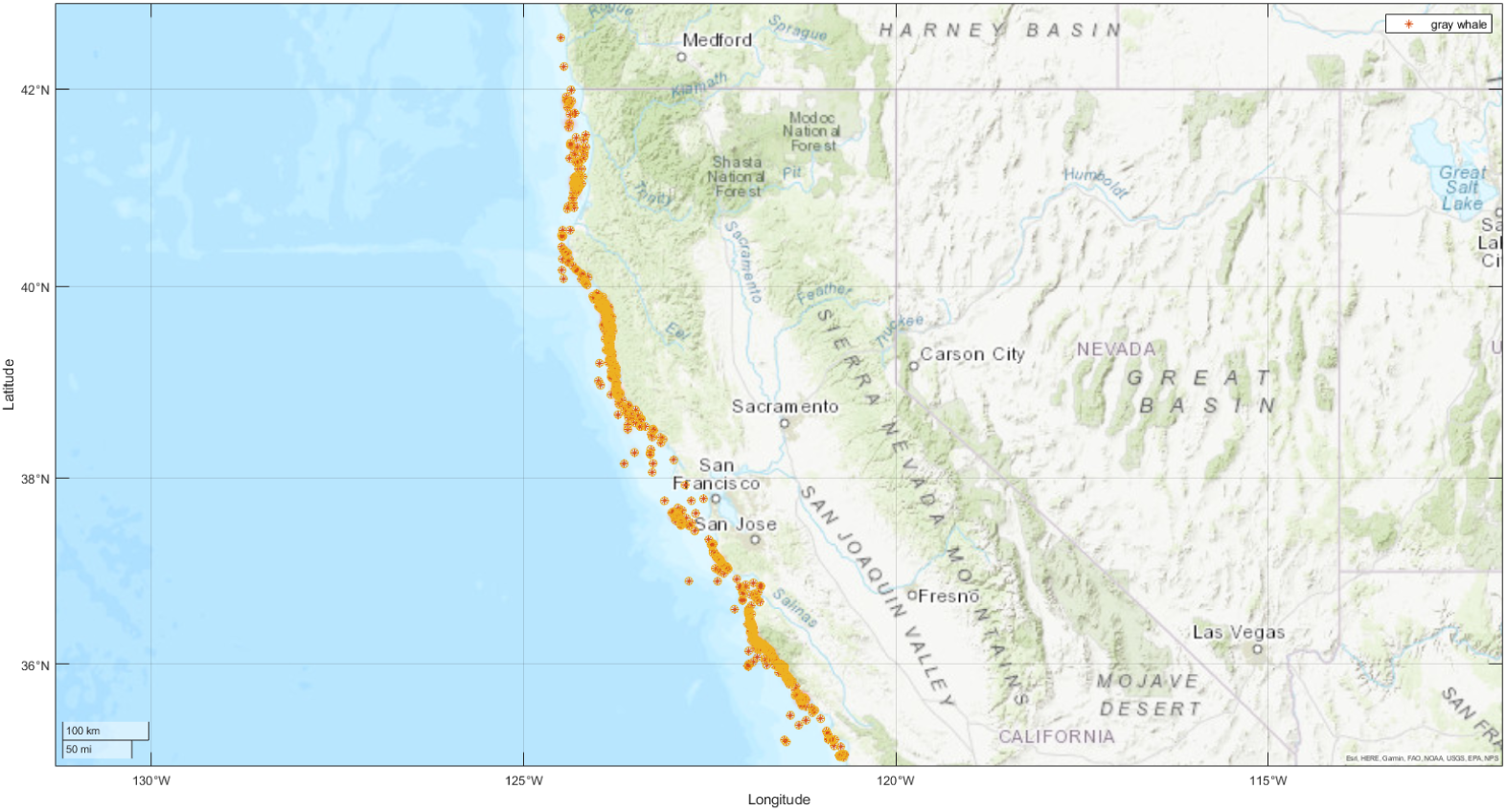
Detections and density map of gray whale detections (northern survey effort) (click image and follow link to zoom in)

**Figure 4.2.4.**
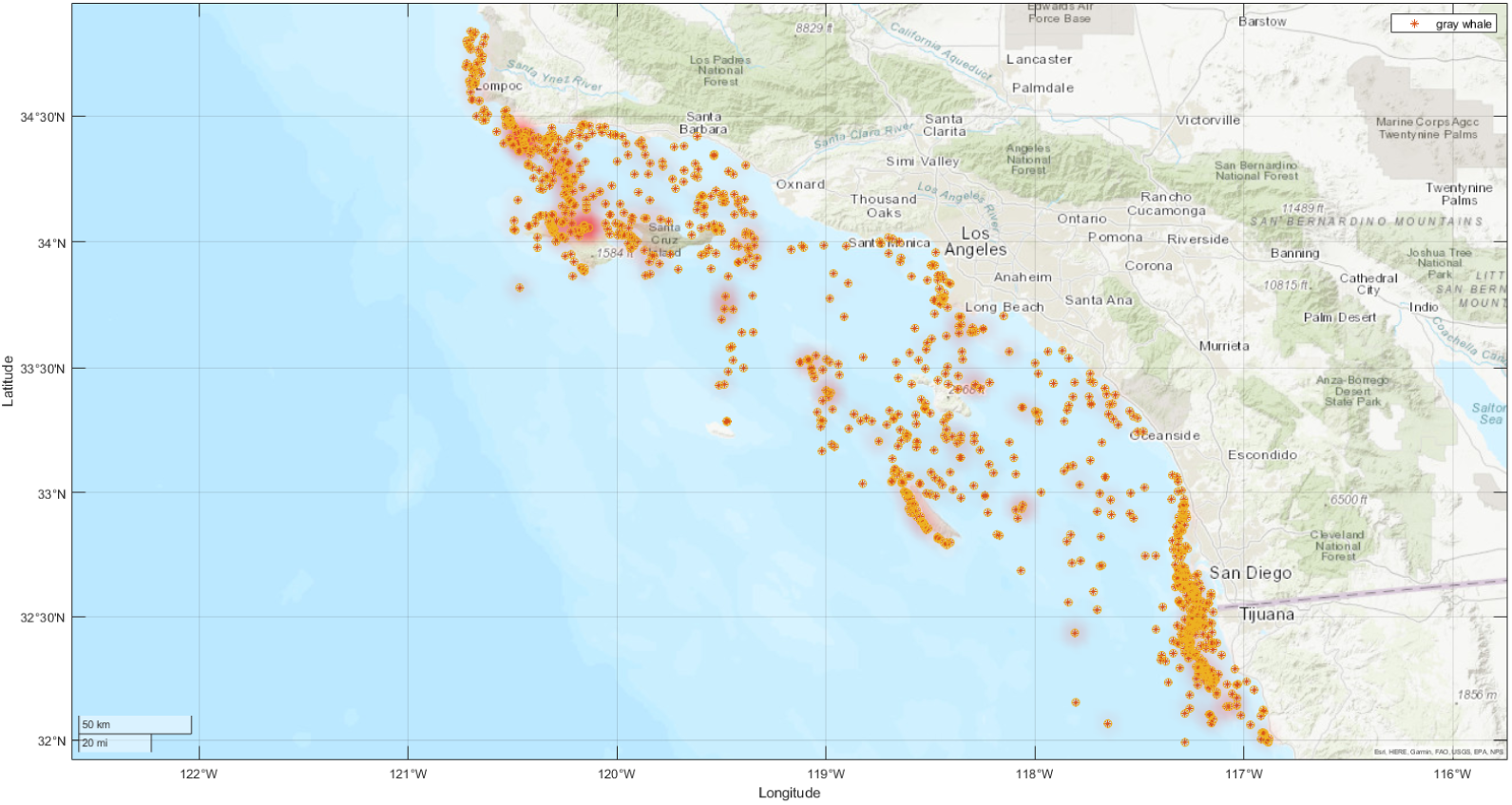
Detections and density map of gray whale detections (southern survey effort)

**Figure 4.2.5.**
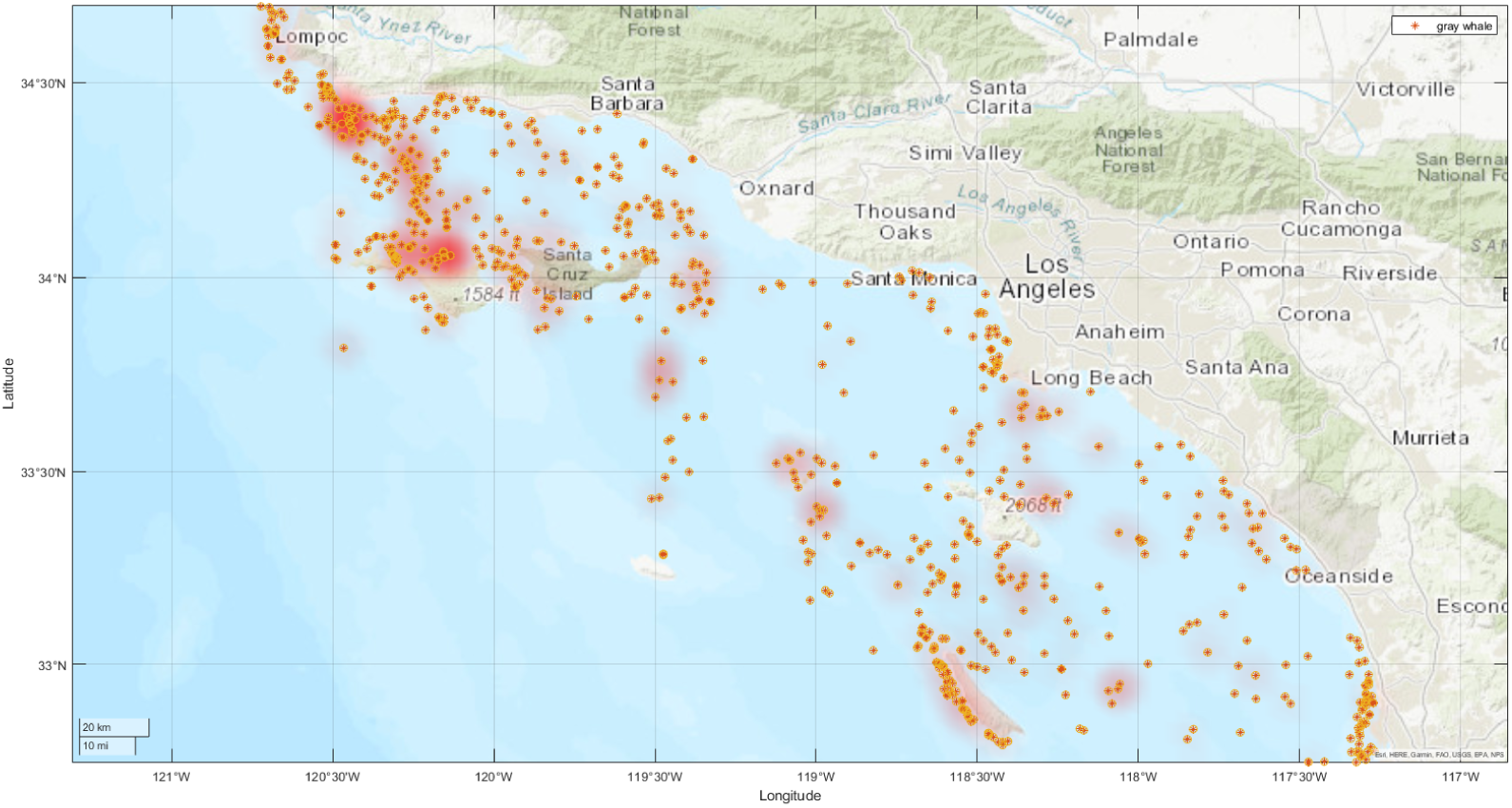
Detections and density map of gray whale detections in the Channel Islands

## Discussion

The detection and classification of large whales in satellite imagery presents a significant technical challenge, particularly in terms of recognition reliability. We are constantly faced with epistemological questions: can whales be consistently recognized? How can we confirm that the detected objects are indeed whales? Can we go further and identify a particular species and discriminate between species? We might then wonder what objects or phenomena actually produce false positives in automated detection pipelines.

The increasing availability of high-resolution satellite imagery, combined with the predictable seasonal presence of certain well-studied species in wintering or migratory grounds, offers a promising foundation for improving large whale detection capabilities. Species such as gray whales, known for their highly predictable migratory behavior and seasonal presence in Baja California’s coastal lagoons, are ideal for developing and validating detection algorithms. Other species with similarly strong site fidelity include southern right whales (*Eubalaena australis*) at locations such as Head of Bight[46], across the Southern Cape coast in South Africa[47] or Peninsula Valdés in Argentina[48], and humpback whales (*Megaptera novaeangliae*) in their Australian wintering grounds [49]. The large scale aggregations provided by those species provide high-confidence opportunities for reliable image annotation, which in turn helps to improve the performance of automated classification systems.

In this study, prior ecological knowledge of gray whale migratory timing [13] was used to build a reliable dataset and to guide the selection of satellite imagery for model training and evaluation. Initial annotation efforts focused on breeding grounds such as Laguna San Ignacio and Bahía Magdalena, where shallow water facilitates visual detection. While turbid conditions introduce biases in color representation, the low probability of other large whale species being present in these lagoons [50] — reduces the likelihood of misclassification. Only bottlenose dolphins are sighted year-round in Ojo de Liebre, San Ignacio and Bahia Magdalena lagoons. Accordingly, detections from these sites were assumed to represent gray whales with high confidence.

One challenge was the identification of calves, which due to their smaller size often fall below detection thresholds. In most cases, smaller masses observed in proximity to or atop adult whales were assumed to be calves. Advances in satellite imaging, such as the availability of sub-15 cm resolution imagery from platforms like Albedo[51], and emerging tools such as super-resolution image reconstruction [52], may significantly improve our ability to resolve such features in future datasets.

It is important to note that while breeding ground observations provide useful priors on whale appearance, substantial variability in sea state, water depth, and spectral reflectance in open-ocean conditions can lead to different visual signatures. While the body shape of surfacing individuals remains largely consistent and can still inform classification, animals at depth may be harder to classify to the species level due to distortion and decreased contrast.

Annotated examples from wintering grounds were used to train models and assess detection validity by comparing size and shape characteristics. However, this approach lacked a formalized similarity metric to quantify resemblance between test samples and reference examples. Developing a quantitative similarity measure remains a crucial next step, potentially using image feature descriptors such as Scale-Invariant Feature Transform (SIFT) [53], or deep learning-based embeddings obtained via pretrained convolutional neural networks or siamese architectures designed for one-shot learning [54].

A potential source of misclassification stems from the visual similarity between gray whales and other large baleen whales, particularly blue (*Balaenoptera musculus*) and fin whales (*Balaenoptera physalus*), which co-occur with gray whales in the California Current ecosystem [9]. While humpback whales are visually distinct, juvenile blue or fin whales may exhibit size and coloration within the range of adult gray whales, increasing the potential for confusion. To our knowledge, training datasets used in this study did not include mislabeled examples of blue or fin whales. Nonetheless, constructing a curated database of these concurrent species is necessary to improve model robustness and reduce false positives. Developing such a reference database is complicated by the relatively low population density and unpredictable spatial-temporal distribution of blue and fin whales, which lack clearly defined wintering areas comparable to gray or humpback whales. Consequently, large-scale image screening and analysis are needed to identify relevant samples.

In addition to the aforementioned visual resemblance of several whale species , environmental factors such as sea state, sun glint, and turbidity have an influence on detection accuracy and may impact similarity estimation. Incorporating multispectral data — including shortwave infrared and near-infrared bands — may enhance classification robustness under varying conditions.

## Conclusions and Future Work

Current spatial planning relies on marine mammal vessel or aerial based surveys, telemetry tags, or opportunistic sightings data which can be biased in space and time. This study presents an important case study towards testing the feasibility of large-scale deployment of a gray whale detection algorithm off of California and more broadly for large-scale whale detection in VHR satellite imagery. Our results demonstrate the feasibility of high accuracy whale detection in VHR satellite imagery as a gray whale monitoring tool in California.

After adequate signal preprocessing, a California gray whale dataset was developed to train a deep-learning-based architecture that uses multispectral and spatial information contained in all the multispectral bands. Under this approach, gray whale detection performance peaks above 99% accuracy, indicating that feature extraction is a viable pathway when a limited amount of data is available, as is frequently the case for whale detection problems.

This performance minimized both false positives and negatives and allows for a rapid assessment of satellite data. In the context of big spatial data, where even low false positive rates produce a number of false positives that can rapidly become intractable for human review, those results are extremely encouraging not only from a theoretical point of view but also for practical use as hinted by a manageable large-scale deployment. Further, even faint images of whales and whales located deeper in the water column that typically result in false negatives were detected. While detection capabilities will continue to improve, current performance is adequate to begin applying this tool to fill information gaps for monitoring and conservation of gray whales off California.

Though the proposed approach is a step towards fully automated whale detection, a significant bottleneck constraining the extension of our California gray whale detection approach to other geographies and whale species is the dependence on the underlying predetection dataset. We note that our results are dependent on the quality of this dataset and its suitability for the area and species of interest. For our study, we opportunistically built a sufficiently sized predetection dataset by targeting known gray whale aggregation sites outside the context of our study area (specifically Baja California, Mexico) to streamline human annotation of whale pixels. The predetection dataset contains whales clearly visible at the ocean surface as well as faint images of whales located deeper in the water column, yet our predetection and training dataset are biased towards detectable whales in relatively low turbidity and low to moderate sea states.

Future work will consist of: (a) assessing the response of the algorithm to higher sea states in terms of false positives and false negatives, which can be achieved by performing a statistical analysis of the deployment datasets presented in section 4.2. and will also involve (b) retraining the algorithm and increasing its performance under a broader range of conditions and the inclusion of a variety of other events that could potentially occur and are currently grouped under the non-whale detectable class (seabirds, dolphins, fishing nets, rocks) as well as other marine mammals). Results will also be made more robust not only by extending the number of classes but also by including more training and testing instances of each class. This is particularly important for the background class which, as it is, already included adverse conditions (wavy surface, fog) but did not include the most adverse weather condition imagery in which whales were seldom observed, that were difficult to manually annotate, and that could potentially trigger false positives. The detector and extracted statistical information already provide a basis to rapidly increase the number of detections which will in turn allow to build and train increasingly robust detectors. Incorporating automated detection of gray whales into our management toolbox can support the analysis of migration timing, the production of reliable abundance estimates (particularly those outside of shore-based sampling approaches), and sampling of undersampled habitats. As migration behavior continues to be shaped by a changing climate, incorporating additional sampling approaches becomes critical in conservation and protection of these marine megafauna.

## Notes

### Competing Interest Statement

The authors have declared no competing interest.

